# Predictive and motivational factors influencing anticipatory contrast: A comparison of contextual and gustatory predictors in food restricted and free-fed rats

**DOI:** 10.1101/2021.05.20.444943

**Authors:** Jessica Hayes, Celia Garau, Giulia Chiacchierini, Gonzalo P. Urcelay, James E. McCutcheon, John Apergis-Schoute

## Abstract

In anticipation of palatable food, rats can learn to restrict consumption of a less rewarding food type resulting in a binge on the preferred food when it is made available. This construct is known as anticipatory negative contrast (ANC) and can help elucidate the processes that underlie binge-like behavior as well as self-control in rodent motivation models. In the current investigation we aimed to shed light on the ability of distinct predictors of a preferred food choice to generate contrast effects and the motivational processes that underlie this behavior. Using a novel set of rewarding solutions, we directly compared contextual and gustatory ANC predictors in both food restricted and free-fed Sprague-Dawley rats. Our results indicate that, despite being food restricted, rats are selective in their eating behavior and show strong contextually-driven ANC and do so after fewer training sessions than free-fed animals. These differences mirrored changes in palatability for the less preferred solution across the different sessions as measured by lick microstructure analysis. Moreover, in contrast to previous research, gustatory cues in both food restricted and free-fed rats were sufficient for ANC to develop although this flavor-driven ANC did not relate to a corresponding change in lick patterning. These differences in the lick microstructure between context- and flavor-driven ANC indicate that the motivational processes underlying ANC generated by the two predictor types are distinct.

## Introduction

In modern society, eating has a time and a place but when faced with strong sensory cues linked to food the temptation to eat can be overwhelming. The sights, sounds and smells of a familiar kitchen while dinner is being prepared can evoke a strong drive to eat but in refraining from doing so one can reserve their appetite for the main course. Such consummatory choice behavior is also at play in foraging animals where decisions based on prior knowledge of territorial food sources can result in animals passing up a nutritionally-insufficient option for one of greater value that is likely available (Stenset et al., 2016, Hertel et al., 2016, and reviewed in Pyke et al., 1977; Timberlake et al., 1987; Kamil & Roitblat, 1985). Such behavior highlights the ability of animals to integrate past information regarding food availability to implement an effective foraging strategy.

The behavioral principles that underlie intertemporal decision-making regarding food have been well studied experimentally across various scientific disciplines (Naik and Moore, 1996; Rosati et al., 2007; Wikenheiser et al., 2013). In one such recent investigation, Billard et al. (2020) observed that when a preferred prey was available at night cuttlefish acted selectively in their food choices, choosing to forgo a less preferred option (crab) in the day and consuming more of their preferred option (shrimp) at night. When shrimp were subsequently made unavailable the same cuttlefish adapted their behavior and became more opportunistic in their feeding choosing to eat more crab during the day. Ultimately, these animals were able to base their prey choices on prior experiences of availability to optimize their feeding behavior.

A related intertemporal choice paradigm has been frequently used for investigating food-related contrast effects in rodents (Flaherty & Largen, 1975; Lucas et aI., 1990; Cottone et al., 2008). Here, in anticipation of a palatable food source rats learn to restrict the consumption of a less rewarding food type, resulting in a binge on the preferred food later in the session when it is made available. These sessions are compared with others in which the preferred food is not made available and the less rewarding food is provided throughout. Over multiple sessions rats reduce their intake of the less preferred food source selectively in sessions where the preferred food follows. This predictive restriction of food intake is called anticipatory negative contrast (ANC) and has been shown to develop with different types of food sources as well as drugs of abuse of different hedonic value (Lucas et aI., 1990; Flaherty and Mitchell, 1999; Grigson and Twining, 2002; Cottone et al., 2008; Parylak et al., 2012) – with the disparity between the two comparative rewards being of utmost importance in developing a contrast effect (Flaherty and Rowan., 1986; Flaherty et al., 1991; Moss et al., 2002). Experimentally, whether or not the session is one where the less preferred food will be followed by a more preferred option is often signposted by sensory information such as discrete sensory stimuli, contextual cues and/or gustatory sensations where solutions are flavored while keeping the nutritional content equivalent (Lucas and Timberlake, 1992; Flaherty et al. 1995). These modulating cues or “occasion setters” are more or less effective in ANC development (Flaherty and Largen, 1975; Lucas and Timberlake, 1992; Wright et al., 2013), indicating that an accurate memory of the predicted reward in a given environment is required for contrasting effects to occur.

While a robust reduction of consumption in anticipation of a future reward is well established using ANC paradigms, the psychological processes underlying this phenomenon are less known. In their 1994 study, Flaherty et al proposed three mechanisms for explaining ANC behavioral manifestations: 1) a progressive devaluation of the first food source when anticipating a preferred option; 2) competing behavioral responses, such as spatial competition with the unavailable port, reducing the amount of time dedicated to licking/eating; and 3) active inhibition of the urge to consume the less preferred option despite animals, in some experiments, being motivated by food restriction. While interpretations based on data from variations of the ANC paradigm mostly support ANC resulting from a reward devaluation of the first food source (Flaherty and Rowan, 1995; Wright et al., 2013; see also discussion Onishi and Xavier, 2011) there have been some conflicting findings as to how the motivational state of the animal (i.e. food restriction) (Flaherty et al., 1991; Weatherly et al., 2005; Wright et al., 2013) and the nature of the predictors (contextual vs flavors) (Lucas and Timberlake, 1992; Flaherty et al. 1995) impact ANC development. Food restriction has been shown both to decrease (Wright et al., 2013) but also increase consumption of the less preferred food option (Flaherty et al., 1991; Weatherly et al., 2005), as increasing food intake in an opportunistic fashion would help meet the animals’ metabolic requirements. Moreover, gustatory cues have resulted in no significant contrast effects (Lucas and Timberlake, 1992; Flaherty et al. 1995), a result that has been interpreted as the flavor acting as a secondary reinforcer for the preferred food source thereby facilitating its consumption (Lucas and Timberlake, 1992; Flaherty et al. 1995; Everitt and Robbins, 2000).

In the current study we aimed to shed light on these conflicting data by comparing contextual and gustatory ANC predictors in both food restricted and free-fed rats. To do so we have used a novel carbohydrate (solution 1; maltodextrin) and condensed milk (solution 2) sequence for further establishing a contrast effect with nutritionally-relevant food sources. Interestingly, different parameters of licking behavior have been related to different motivational processes (Davis and Smith, 1992; reviewed in Dwyer, 2012; Johnson, 2018 and Naneix et al., 2019) and as such we have analyzed the lick microstructure to relate changes in lick parameters to changes in the hedonic value of the solutions, as this could shed light on the underlying psychological mechanisms of ANC.

Our results indicate that, despite being hungry and potentially benefiting from opportunistic feeding, rats that are food restricted are selective in their eating behavior and show strong contextually-driven ANC. In addition, they do so after fewer training sessions than their free-fed counterparts. These differences mirrored changes in palatability for the less preferred solution across the different sessions as gauged by lick microstructure analysis. Moreover, in contrast to previous research, gustatory predictive cues in both food restricted and free-fed rats were sufficient for ANC development, an effect that could not be explained by hedonic changes as determined by lick measurements of palatability and a pre/post-conditioning flavor preference test.

## Methods

### Animals

For all experiments 40 male Sprague Dawley rats, of approximately 300 g at the start of testing, were purchased from Charles River (Cambridge, UK) and housed in pairs. The room was kept at 21° C and humidity of between 40% and 70% under a 12 hour light-dark cycle (lights on at 7:00). Rats had free access to standard lab chow (Teklad Global Diet, Envigo) and water. Prior to the experiment half of the rats were put on a mild food restriction, and given 12.5 g of lab chow per day. This food restriction continued on experimental days but rats were put on a free-access diet during the weekends. All rats were weighed daily Monday through Friday and a percentage of baseline weights calculated. All testing was conducted in accordance to the Animals (Scientific Procedures) Act 1986 (PPL# PFACC16E2).

### Behavioural Protocol

Animals were trained and tested in two identical operant chambers (30.5 × 24.1 × 21.0 cm; Med Associates), each located inside a sound- and light-attenuated aluminum outer chamber (1200 × 700 × 700 cm). The behavioral chambers were equipped with a house light located on the left wall and 2 retractable sippers located on the right wall, which when extended fully were located approximately 1 cm behind the chamber wall. Sippers were accessed via ports (oval shaped H: 1.5 cm W: 1.1cm) in the wall through which rats needed to poke their noses. This arrangement ensured that only the tongue could contact the sipper and prevented the formation of fluid bridges meaning that individual licks were recorded with high fidelity. Ports were also fitted with infrared beam breaks for detecting port entries. Contact lickometers (Med Associates) were used to detect number of licks for each solution. The house light was turned on at the beginning of each daily session and turned off at the end of it. Equipment was controlled by a computer running Med-PC IV Software Suite (Med Associates). Webcams were used to monitor the animals’ behavior.

Animals were pre-trained for 10 min per day to lick a palatable 10% sucrose solution in all experimental boxes (Context A and B) for 6 days prior to the experiment. For experiments using a contextual predictor, a standard Med Associates box with clear plexiglass walls and a barred floor was used for Context A and a modified version with a striped wall covering, a fine wire mesh floor and continuous white noise (75 dB level) played on a speaker was used for Context B. Both sippers were extended simultaneously for 5 minutes, retracted for 20 sec and presented again for a further 5 minutes. Licks were recorded from both sippers at this time to check that there was no location bias within the contextual behavioral paradigm.

The ANC conditioning protocol (Figure 1) consisted of one session run across two days, a control and an experimental day. On Day 1 (control day) of context predictor experiments, sipper 1 containing a 2% maltodextrin + 0.2% sodium saccharin (Malt) solution was extended (phase 1; 5 min) followed by an inter-phase interval (IPI; 20 s) during which both sippers were retracted and finally extension of sipper 2 containing an identical solution as phase 1 (phase 2; 5 min). On Day 2 of each session (experimental day) of context predictor experiments, animals were placed in the second context and sipper 1 was extended containing Malt solution, identical to control day (phase 1; 5 min). This was followed by an IPI (20 s) with no sippers extended and finally, sipper 2 extended containing a 50% Malt/condensed milk (CM) solution (phase 2; 5 min). For flavor predictor experiments, the context was always identical and flavor was used as a predictor by adding either grape or cherry Kool-Aid (0.05%) to solutions. Specifically, flavor A was added to both phase 1 and phase 2 solutions on control days (phase 1, Malt / phase 2, Malt) whereas flavor B was added to both solutions on experimental days (phase 1, Malt / phase 2, CM). For all ANC conditioning experiments the first sipper location (left or right) and the predictor (i.e. flavor and context) were counter-balanced. Number of licks, number of head entries, number of lick clusters and licks per cluster were recorded for each session, and animal weight was recorded daily. At the end of the experiments animals were humanely culled via a schedule 1 method.

**Figure 1:**
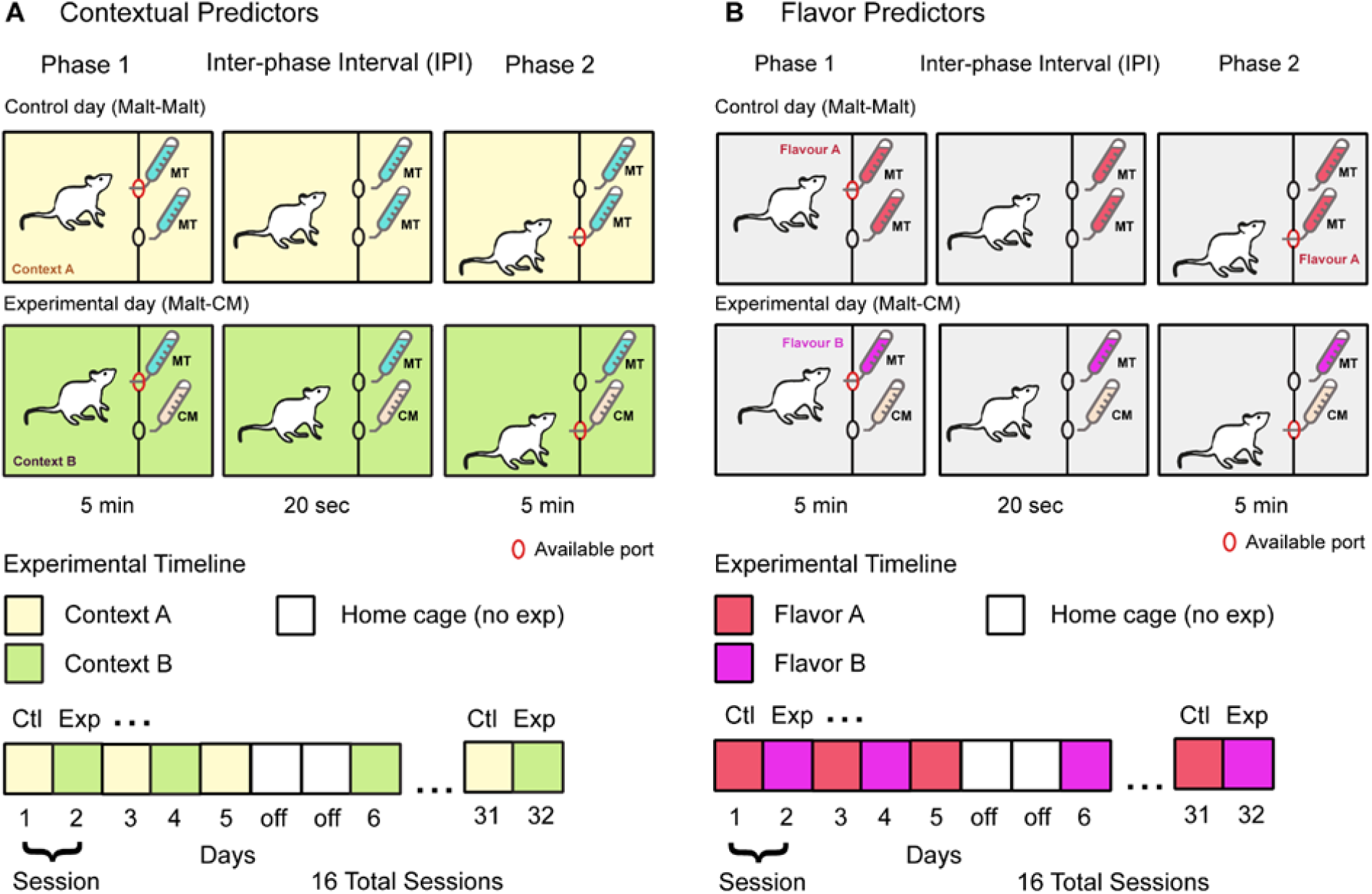
Anticipatory Negative Contrast paradigm. **A**. Using contextual cues as predictors, on alternate days a different context predicted either a condensed milk (CM) or the same maltodextrin (Malt) solution in phase 2 (5 min) as the one given in phase 1 (5 min). An interphase interval (IPI) where no sipper was extended separated the two phases (20 sec). **B**. Using flavored maltodextrin solutions in phase 1, on alternative days a different flavored solution predicted either a condensed milk or the same maltodextrin solution in phase 2 (5 min) as the one given in phase 1 (5 min). An interphase interval (IPI) where no sipper was extended separated the two phases (20 sec).

For the flavor preference test performed before and after ANC conditioning, two sippers each containing one of two flavors (grape or cherry) + 0.2% sodium saccharin were extended one at a time for a maximum time of 30 seconds, or 10 seconds after first sipper contact. Overall, 10 “forced choice trials” where only 1 sipper was presented at a time was followed by 30 “free choice trials” where both sippers were presented together.

### Statistical Analysis and Code Availability

Behavioral data (lick and port entry timestamps) were extracted from data files and analyzed using custom Python scripts that measured numbers of licks for each solution (DOI: 10.5281/zenodo.4772860). Lick microstructure was analyzed by using interlick intervals to divide licks into clusters (Davis and Smith, 1992). Clusters were defined as runs of licks with no interlick intervals > 500 ms. For statistical analysis of within session behavioral variables, two-way repeated measures ANOVA were used with Condition (control vs. experimental) and Session as within subject variables. For two-way statistical analyses compared Condition (control vs experimental) and Session (ANC conditioning days) and for three-way ANOVA, Diet (FF and FR) was included as a variable.

## Results

### Experiment 1: Anticipatory Negative Contrast Using Contextual Predictors

#### Total licks: Contextual predictors drive ANC in free-fed and food restricted rats

We used a modified ANC paradigm with a novel sequence of rewarding solutions to gauge the effectiveness of contextual or gustatory predictors for enhancing contrast effects in free fed (FF) and food restricted (FR) Sprague-Dawley rats (Figure 1). When using contextual cues as predictors, we saw clear ANC develop in FF rats (n = 10) as evidenced by reduced consumption during phase 1 specifically on days when a more preferred solution was expected. Statistically, total lick measurements revealed a significant effect by ANC conditioning day (session) (2-way ANOVA: F(1,9)=6.20, P=0.03) and interaction between session and Malt-Malt and Malt-CM days (condition) (F(7,63)=4.62, P=0.0003) (Figure 2A1-left). Post-hoc Bonferroni analysis showed significant differences in total phase 1 licks between Malt-Malt and Malt-CM days on sessions 6, 7 and 8. As expected in phase 2, FF rats (Figure 2A1-right) increased their lick rates to the highly palatable CM solution on Malt-CM days compared to the less preferred phase 2 Malt solution on Malt-Malt days (FF, 2-way ANOVA, Condition: F(7,63)=8.46, P<0.0001; Session: F(1,9)=36.70, P=0.0002; Interaction: F(7,63)=9.17, P<0.0001; Bonferroni post-hoc differences - Sessions 2 through 8). Using a novel sequence of rewards in the current study, these results extend previous findings that FF rats can develop robust ANC (Flaherty et al., 1991; Cottone et al., 2008, Giuliano et al., 2012).

**Figure 2:**
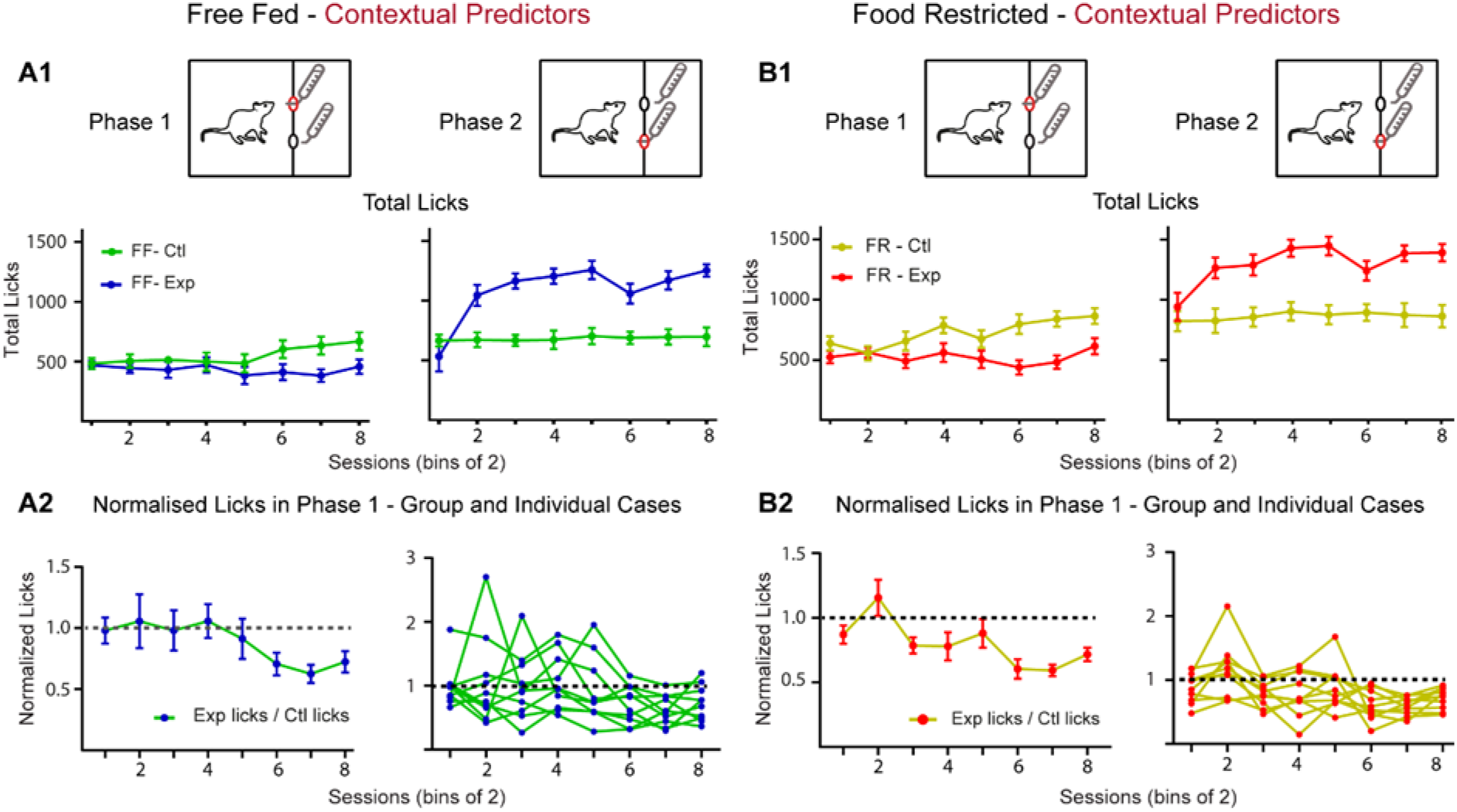
Using contextual predictors, free-fed and food restricted animals develop negative contrast. **A1**. For ad libitum fed animals, in phase 1 a negative contrast in total licks for a maltodextrin solution develops over paired Malt-Malt/Malt-CM sessions (left). In phase 2, CM consumption increases early on in training and is maintained throughout the experiment (right). **A2**. Data showing normalized lick data for paired Malt-Malt and Malt-CM sessions as a group (left) and for individual cases (right). **B1**. For food restricted fed animals, in phase 1 a negative contrast in total licks develops over paired Malt-Malt/ Malt-CM sessions (left). In phase 2, CM consumption increases early on in training and is maintained throughout the experiment (right). **B2**. Data showing normalized lick data for paired Malt-Malt and Malt-CM sessions as a group (left) and for individual cases (right). Group analyses show that contrast effects develop faster in FR compared to FF animals (**A2**,**B2** – left panel). FF, free-fed; FR, food restricted.

FR rats (n=10) also showed strong ANC (Figure 2B1-left). Statistically, a significant effect was seen by condition (Malt-Malt vs. Malt-CM: 2-way ANOVA: F(7,63)=3.94, P=0.001) and by conditioning day (Session: (F(1,9)=21.94, P=0.001) and an interaction between the two was also revealed (F(7,63)=6.03, P<0.0001). These results are consistent with a recent within-subject ANC demonstrating that contextual cues are sufficient for ANC to develop in FR rats, albeit using a different sequence of rewards (Wright et al. 2013). Interestingly, post-hoc Bonferroni tests showed significant differences between control and experimental conditions on sessions 3 through 8 indicating that in FR animals ANC developed after fewer conditioning trials than FF rats. As expected in phase 2, FR animals increased their licking to CM compared to Malt (2-way ANOVA, Group: F(7,63)=3.94, P=0.0013; Sessions: F(1,9)=21.94, P=0.001; Interaction: F(7,63)=6.03, P<0.0001; Bonferonni post-hoc differences –Sessions 3 through 8). Overall, these results indicate that contextual cues can act as effective signals for decreasing animals’ intake of a less palatable Malt solution in anticipation of a more preferred CM option whose intake increases when it is made available. Moreover, FR animals developed ANC after fewer conditioning trials (session 3) than FF animals (session 6) suggesting that despite being hungry FR can act selectively in their feeding behavior.

#### Premature port entries: Contextual predictors drive ANC in free-fed and food restricted rats

A commonly used ANC measure is based on the amount of sipper contact during licking. In addition to quantifying contrast effects using total licks, we measured entries to the phase 2 sipper port as a second measure of anticipation (Figure 3). In addition, this analysis aims to shed light on the contribution of competing behavioral responses (i.e. spatial competition with the unavailable sipper) in reducing the amount of time spent licking/eating that may contribute to a decrease in total licks on Malt-CM days. Differences in premature port entries between Malt-Malt and Malt-CM days were seen in both FF and FR rats (FF: 2-way ANOVA, Condition: F(1 9)=19.91, P=0.002; Session: F(7,63)=6.54, P<0.0001; Interaction: F(7,63)=10.05, P<0.0001; Bonferroni post-hoc differences –Sessions 3,4,6-8) (Figure 3A1) (FR: 2-way ANOVA, Condition: F(1,9)=29.59, P=0.0004; Interaction: F(7,63)=4.16, P=0.0008; Bonferroni post-hoc differences –Sessions; 3-8; No main effect by Session: F(7,63)=1.88, P=0.087) (Figure 3B1) further indicating that contextual cues can act as effective predictors for ANC development. Notably, the first of these significant differences by session either preceded or occurred at the same time as the differences in total licks (FF: Total Licks, Session 6; Premature entries, Session 3; FR: Total Licks, Session 3, Premature Port Entries, Session 3). Moreover, the magnitude of premature port entries did not closely match ANC levels (Figures 2, 3) overall suggesting that spatial competition for the unavailable sipper cannot entirely account for the reduction in total licks on Malt-CM days.

**Figure 3:**
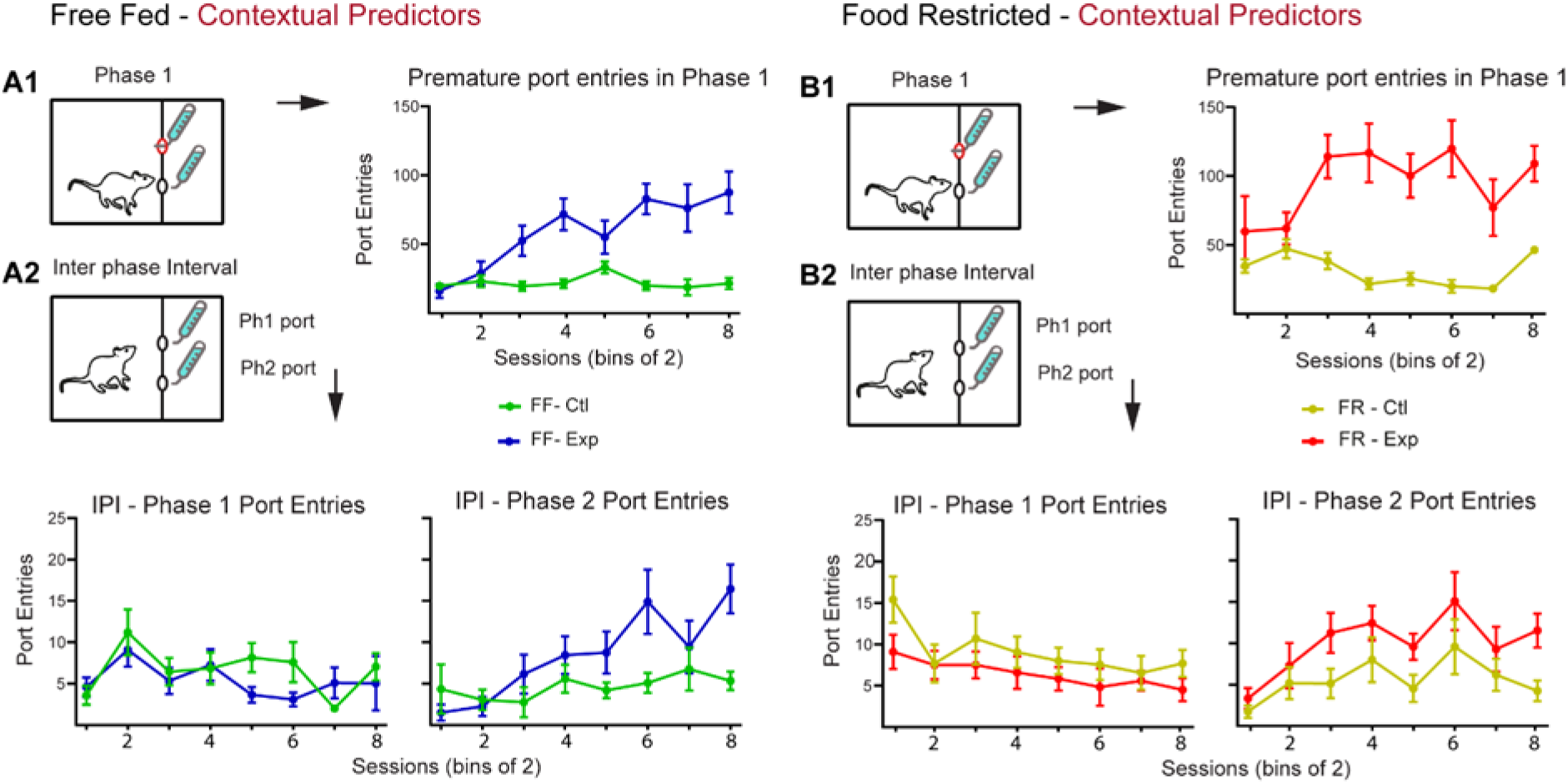
Premature port entries using contextual contrast cues. **A1**. On Malt-CM days, FF animals **(A)** made significantly more premature port entries in phase 1. **A2**. During the IPI animals on Malt-CM days progressively made more premature phase 2 port entries than on Malt-Malt days. **B1**. On Malt-CM, FR animals **(B)** made significantly more premature port entries in phase 1than they did on Malt-Malt days. **B2**. During the IPI animals on Malt-CM days progressively made more premature phase 2 port entries than on Malt-Malt days (right). There was no difference in phase 2 port entries between Malt-Malt and Malt-CM sessions in either FF or FR animals. FF, free-fed; FR, food restricted.

These premature entry results are in line with data collected during the IPI phase where port entries to unavailable sippers 1 and 2 was measured for 20 sec (Figure 1-IPI). On Malt-CM days, both FF (Figure 3A2-right) and FR (figure 3B2-right) animals made more premature entries into the port where CM availability was anticipated (FF, 2-way ANOVA, Condition: F(1,9)=12.62, P=0.006; Sessions: F(7,63)=4.39, P=0.0005; Interaction: F(7,63)=3.36, P=0.0041; Bonferroni post-hoc differences – Sessions 6,8) (FR, 2-way ANOVA, Condition: F(1,9)=11.64, P=0.008; Session: F(7,63)=3.84, P=0.002; Bonferroni post-hoc differences –Session 8; No interaction: F (7,63) = 0.66, P=0.70). No differences for port 1 entries were seen in any condition (FF, 2-way ANOVA, Condition: F(1,9)=1.20, P=0.30; Session: F(7,63)=2.87, P=0.05; Interaction: F(7,63)=1.32, P=0.29) (Figure 3A2-left) (FR Condition: F(1,9)=4.50, P=0.06; Paired Session: F(7,63)=2.27, P=0.05; Interaction: F(7,63)=0.58, P=0.77) (Figure 3B2-left). Overall, using premature port2 entries as a second measure of anticipation these results further show that contextual cues are sufficient predictors for robust ANC development.

#### Lick microstructure analysis reveals hedonic changes in reward properties related to ANC

To shed light on the hedonic factors that may underlie context-driven ANC we analyzed the lick microstructure of FF and FR rats during all phases and conditions. The amount of continuous licks per cluster - defined by a temporal gap in lick frequency (see methods) - is a measure commonly used to quantify the palatability of the reward (liking) while total clusters has been shown to relate to a reward’s incentive value (wanting) as well as signal its post-ingestive feedback properties (Davis and Smith, 1992; Berridge et al., 2009; reviewed in Dwyer, 2012; Johnson, 2018 and Naneix et al., 2019). Phase 1 licks per cluster analysis for both FF and FR rats showed that a significant difference between Malt-Malt and Malt-CM days developed over time (FF: 2-way ANOVA: F(1,9)=10.87, P=0.009; Session (F(7,63)=2.43, P=0.03; Interaction: (F(7,63)=3.72, P=0.002; Bonferroni post-hoc differences - Sessions 6,7) (Figure 4A1-left). (FR: 2-way ANOVA: Interaction: F(7,63)=3.15, P=0.006; Bonferroni post-hoc difference –Session 6) (Figure 4B1-left). In phase 2, FF animals on Malt-CM days similarly showed a significant increase in licks per cluster compared to Malt-Malt days (2-way ANOVA, Session: F(7, 63)=2.24, P=0.04; Interaction: F(7,63)=4.46, P=0.0004; Bonferroni post-hoc differences –Sessions 1,5,7) (Figure 4A1-right) and so did FR animals (2-way ANOVA, Session: F(1,9)=2.47, P=0.03; Interaction: F(7,63)=2.34, P=0.03; Bonferroni post-hoc differences –Session 3) (Figure 4B1-right).

**Figure 4:**
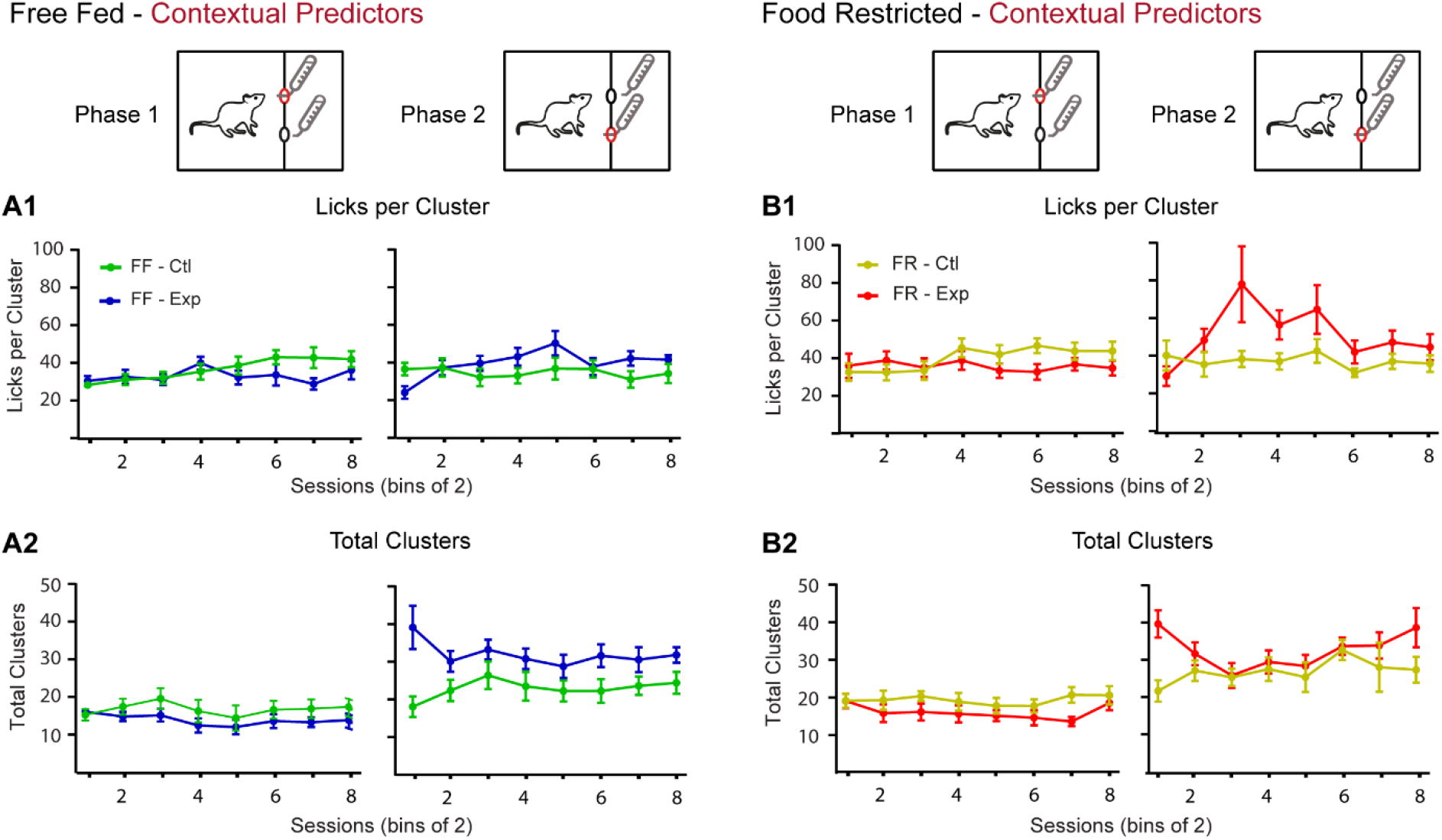
Lick microstructure analysis for contextual contrast experiment. **A1**. In FF animals, differences in amount of licks per cluster progressively emerge between Malt-Malt and Malt-CM sessions in phases 1 (left) and 2 (right). **A2**. In phase 1 on Malt-CM days, FF animals engage fewer lick cluster bouts than on Malt-Malt days (left) while in phase 2 there are more lick cluster bouts compared to Malt-Malt days (right). **B2**. A similar profile is seen in FR animals for licks per cluster **(B1)** and total clusters **(B2)**. FF, free-fed; FR, food restricted.

When total clusters was analyzed no significant differences for either FF or FR animals on Malt-Malt compared to Malt-CM days in phase 1 were seen (FF, 2-way ANOVA, Condition: F(1,9)=2.45, P=0.15; Session: F(7,63)=1.13, P=0.36; Interaction: F(7,63)=0.76, P=0.62) (Figure4A2-left) (FR, 2-way ANOVA, Condition: F(1,9)=3.91, P=0.08; Sessions: F(7,63)=1.38, P=0.23; Interaction: F(7, 63) = 1.02, P=0.42) (Figure 4B2-left). In phase 2 however, there was a significant main effect by Condition and an Interaction between Condition and Session for FF animals (2-way ANOVA, Condition: F(1,9)=8.52, P=0.02; Interaction: F(7,63)=2.82, P=0.01; Bonferroni post-hoc differences – Paired Sessions 1,6; No main effect by Session: F(7,63)=0.72, P=0.65) (Figure 4A2-right) while no effect was seen for FR animals (2-way ANOVA, Condition: F(1,9) = 3.47, P=0.10; Session: F(7,63)=1.42, P=0.21; Interaction: F(7,63)=1.79, P=0.10) (Figure 4B2-right). Overall, these results indicate that, in contrast to total clusters, as ANC develops so do differences in licks per cluster for the Malt solution between Malt-CM and Malt-Malt days.

In summary, when using contextual predictors, ANC was seen to develop in both FF and FR rats and this may be associated with a change in palatability of the non-preferred maltodextrin solution on experimental vs. control sessions.

### Experiment 2: Anticipatory Negative Contrast Using Gustatory Predictors

#### Total licks: gustatory predictors drive ANC in free-fed and food restricted rats

We next tested whether gustatory cues can be used as effective predictors for ANC to develop. In contrast to previous results (Lucas and Timberlake, 1992; Flaherty et al. 1995), FF (n=10) and FR (n=10) rats showed ANC when different flavoring was added to phase 1 Malt solutions used to predict either Malt or CM in phase 2. Specifically, in FF rats (Figure 5A1-left), consumption of Malt in phase 1 was reduced on days in which rats were due to receive CM in phase 2 (2-way ANOVA, Condition: F(1,9)=6.26, P=0.03; Session: F(7,63) = 3.99, P=0.001; Bonferroni post-hoc differences –Sessions 4,5; No Interaction: F(7,63)=1.28, P=0.27). Similarly in FR rats (Figure 5B1-left) the same pattern of reduced phase 1 consumption was seen (2-way ANOVA, Condition: F(1,9)=7.47, P=0.02; Session: F(7,63)=4.26, P=0.0007; Interaction: F(7,63)=3.53, P=0.0029) and was apparent on sessions 4, 6-8 (Bonferroni post-hoc differences). As expected, phase 2 consumption was greater for both groups (Figure 5A1, B1-right on days when CM was available (FF: 2-way ANOVA, Condition: F(1,9)=63.98, P<0.0001; Bonferroni post-hoc differences – Paired Sessions 1-8; No main effect by Session: F(7,63)=0.69, P=0.68; No Interaction: F(7,63)=1.31, P=0.26) (FR: 2-way ANOVA, Session: F(7,63)=2.91, P=0.01; Interaction: F(7,63)=7.15, P<0.0001; Bonferroni post-hoc differences – Sessions 1-4; No effect by Condition: F(1,9)=5.06, P=0.05). In summary these results demonstrate that when Malt is followed by CM gustatory cues are sufficient predictors for ANC development.

**Figure 5:**
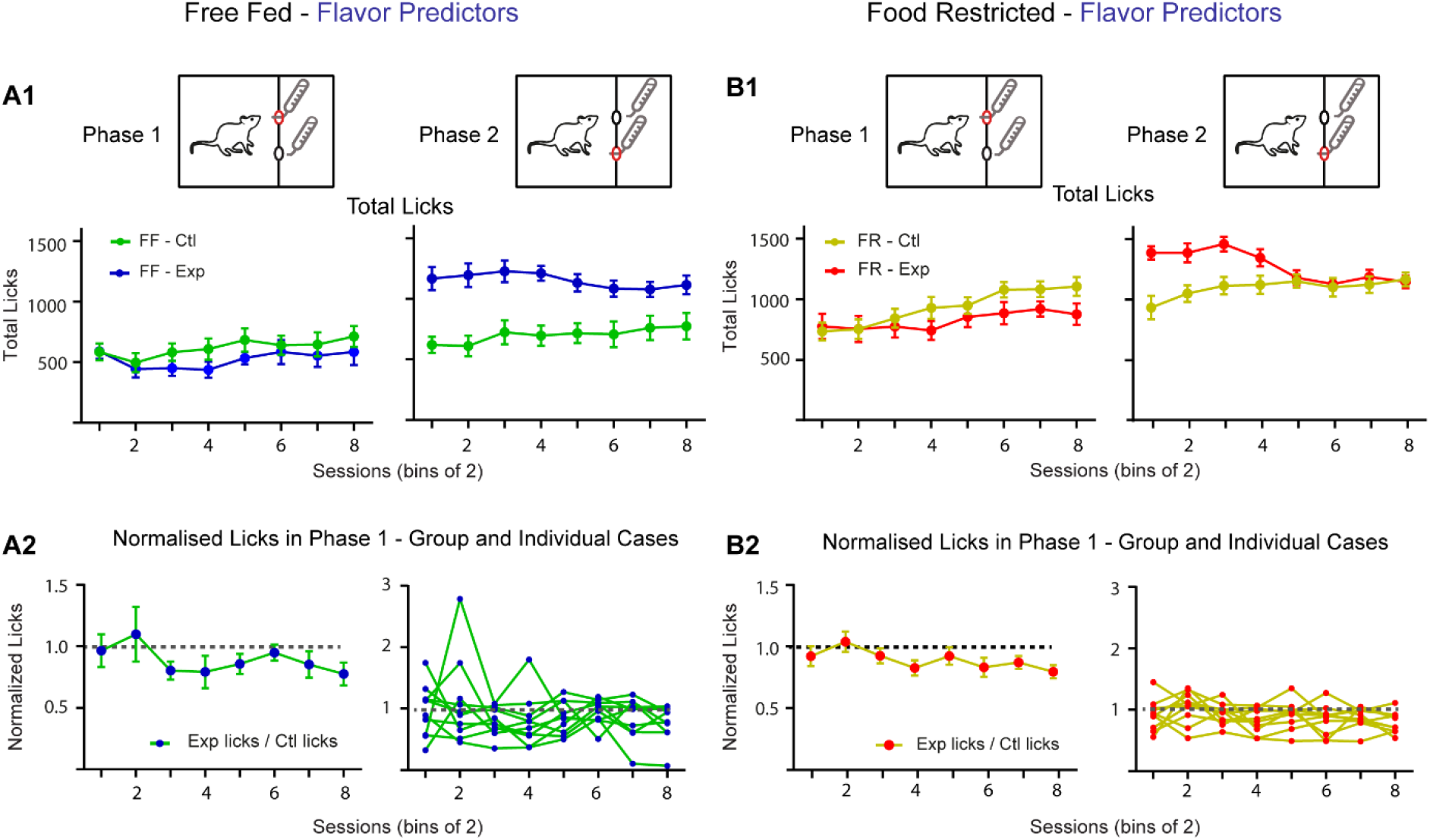
Using flavor predictors, ad libitum fed and food restricted animals develop negative contrast. **A1**. For ad libitum fed animals, in phase 1 a negative contrast in total licks for Malt develops over paired Malt-Malt/Malt-CM sessions (left). In phase 2, CM consumption in FF animals is elevated throughout (right). **A2**. Data showing normalized lick data for paired Malt-Malt and Malt-CM sessions as a group (left), for individual cases (middle) and averaged for each animal for the first and second half of the training (right). **B1**. For food restricted animals, in phase 1 a negative contrast in total licks develops over paired Malt-Malt/ Malt-CM sessions (left). In phase 2, CM consumption is high early in training but decreases to Malt consumption levels towards the end of training (right). **B2**. Data showing normalized lick data for paired Malt-Malt and Malt-CM sessions as a group (left), for individual cases (middle) and averaged for each animal for the first and second half of the training (right). FF, free-fed; FR, food restricted.

#### Premature port entries: gustatory predictors drive ANC in free-fed and food restricted rats

To further support the impression that gustatory cues can act as effective predictors for ANC to develop we next looked at premature phase 2 port entries. In line with the total lick data, both FF and FR rats on Malt-CM days made significantly more premature phase 2 port entries than on Malt-Malt days (FF, 2-way ANOVA, Condition: F(1,9)=31.47, P=0.0003; Session: F(7,63)=2.22, P=0.04; Interaction: F(7 63)=3.70, P=0.002; Bonferroni post-hoc differences –Sessions 2-8) (Figure 6A1) (FR, 2-way ANOVA, Condition: F(1,9)=37.31, P=0.0002; Session: F(7,63)=6.50, P<0.0001; Interaction: F(7,63)=4.77, P=0.0002; Bonferroni post-hoc differences – Paired Sessions 2-8) (figure 6B1). Notably, the first of these significant differences by session preceded significant differences in total licks (FF: Total Licks, Session 4; Premature entries, Session 2; FR: Total Licks, Session 4, Premature Port Entries, Session 2).

**Figure 6:**
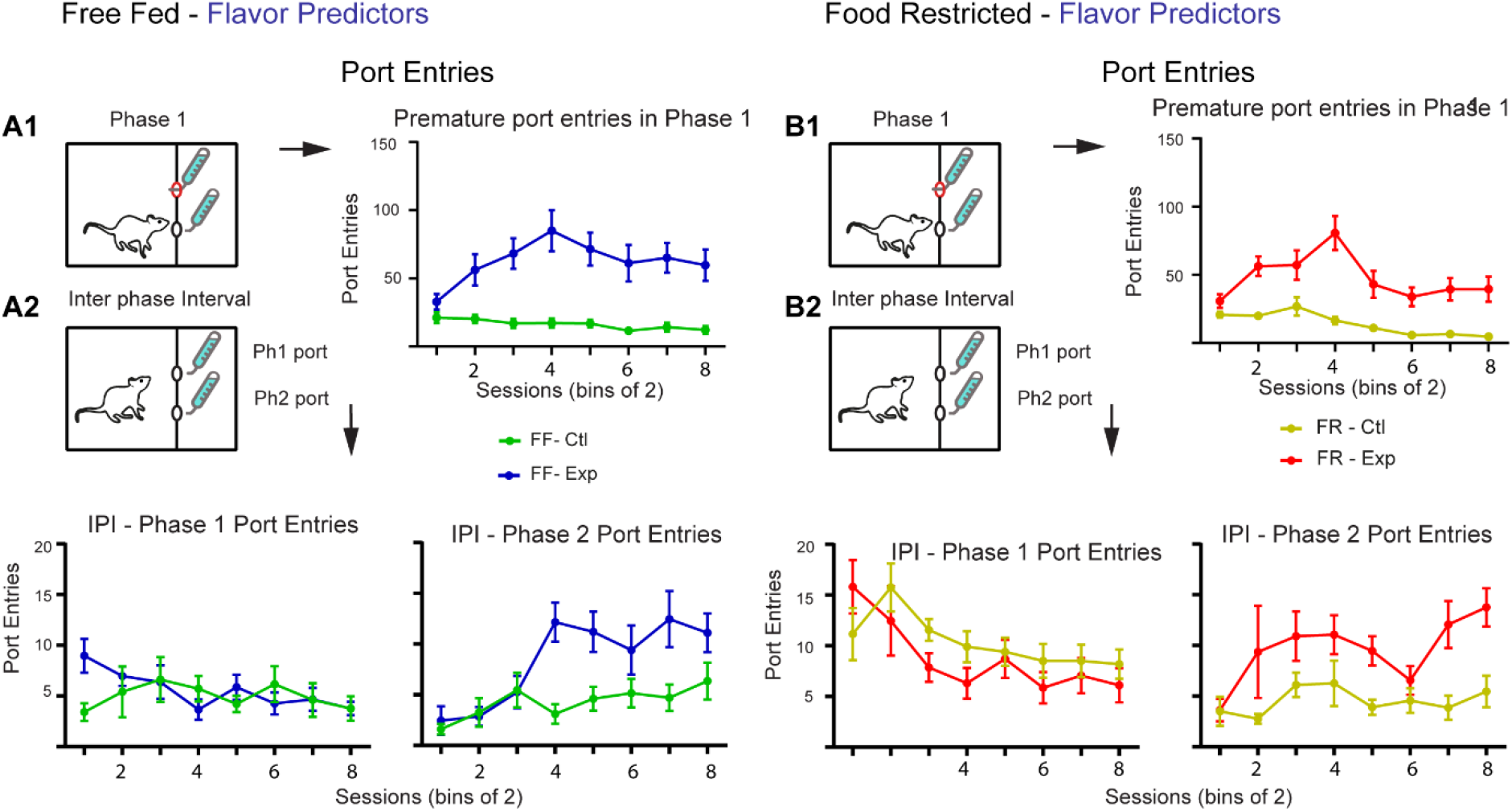
Anticipatory port entries using flavor contrast cues. **A1**. On Malt-CM days, FF animals **(A)** made significantly more premature port entries in phase 1. **A2**. During the IPI, animals on Malt-CM days progressively made more premature phase 2 port entries than on Malt-Malt days. There was no difference in phase 2 port entries between Malt-Malt and Malt-CM sessions (left) during the IPI. **B1**. On Malt-CM, FR animals **(B)** made significantly more premature port entries in phase 1 than they did on Malt-Malt days. **B2**. During the IPI animals on Malt-CM days progressively made more premature phase 2 port entries than on Malt-Malt days (right). FF, free-fed; FR, food restricted.

Similarly, during the IPI both FF and FR animals entered the phase 2 port more frequently during Malt-CM compared to Malt-Malt sessions (FF, 2-way ANOVA, Condition: F(1,9)=13.09, P=0.006; Session: F(7,63)=4.71, P=0.0003; Interaction: F(7,63)=3.39, P=0.004; Bonferroni post-hoc differences – Sessions 4,5,7) (Figure 6A2-right) (FR, 2-way ANOVA, Condition: F(1,9)=21.96, P=0.001; Session: F(7,63)=2.50, P=0.02; Bonferroni post-hoc differences – Sessions 7,8; No significant Interaction: F(7,63)=1.26, P=0.28) (Figure 6B2-right). There was, however, no significant difference between IPI phase 1 port entries on experimental compared to control days for either FF or FR animals (FF, 2-way ANOVA, Condition: F(1,9)=0.78, P=0.40; Session: F(7,63)=0.94, P=0.48; Interaction: F(7,63)=1.56, P=0.17) (Figure 6A2-left) (FR, 2-way ANOVA, Condition: F(1,9)=3.16, P=0.11; Session: F(7,63)=5.47, P<0.0001; Interaction: F(7,63)=1.09, P=0.38) (Figure 6B2-left). These results add support to the data on total licks showing that gustatory cues are sufficient predictors for ANC to develop.

#### Lick microstructure analysis for ANC driven by gustatory cues reveals no change in lick patterning

To determine the underlying hedonic processes behind the total lick changes resulting from using flavored cues we next analyzed the lick microstructure of lick responses influenced by gustatory predictors. In phase 1, there was no difference in licks per cluster for either FF or FR animals on Malt-CM compared to Malt-Malt days (FF: 2-way ANOVA, Condition: F(1,9)=3.29, P=0.10; Interaction, F(7,63)=0.77, P=0.61; main effect by Sessions: F(7,63)=4.42, P=0.0005) (FR: 2-way ANOVA, Interaction: F(7,63)=2.21, P=0.05; Session: F (7,63)=4.0, P=0.001; Bonferroni post-hoc differences – Session 8; No main effect by Condition: F(1,9)=1.54, P=0.24). In phase 2, both FF and FR animals showed a session-dependent change in licks per cluster for CM compared to Malt (FF, 2-way ANOVA, Interaction: F(7,63)=2.88, P=0.01; No main effect by Condition: F(1,9)=2.69, P=0.14; or by Session: F(7,63)=1.68, P=0.13) (FR, 2-way ANOVA, Interaction: F(7,63)=3.63, P=0.002; Bonferroni post-hoc differences – Paired Session 5,8; No main effect by Condition: F(1,9)=4.83, P=0.06 or by Paired Session: F(7,63)=1.04 P=0.41). Notably, these results show that in either motivational state (FF or FR) licks per cluster in phase 2 shifted from high to low levels with conditioning trials (Figure 7A1,B1-right).

**Figure 7:**
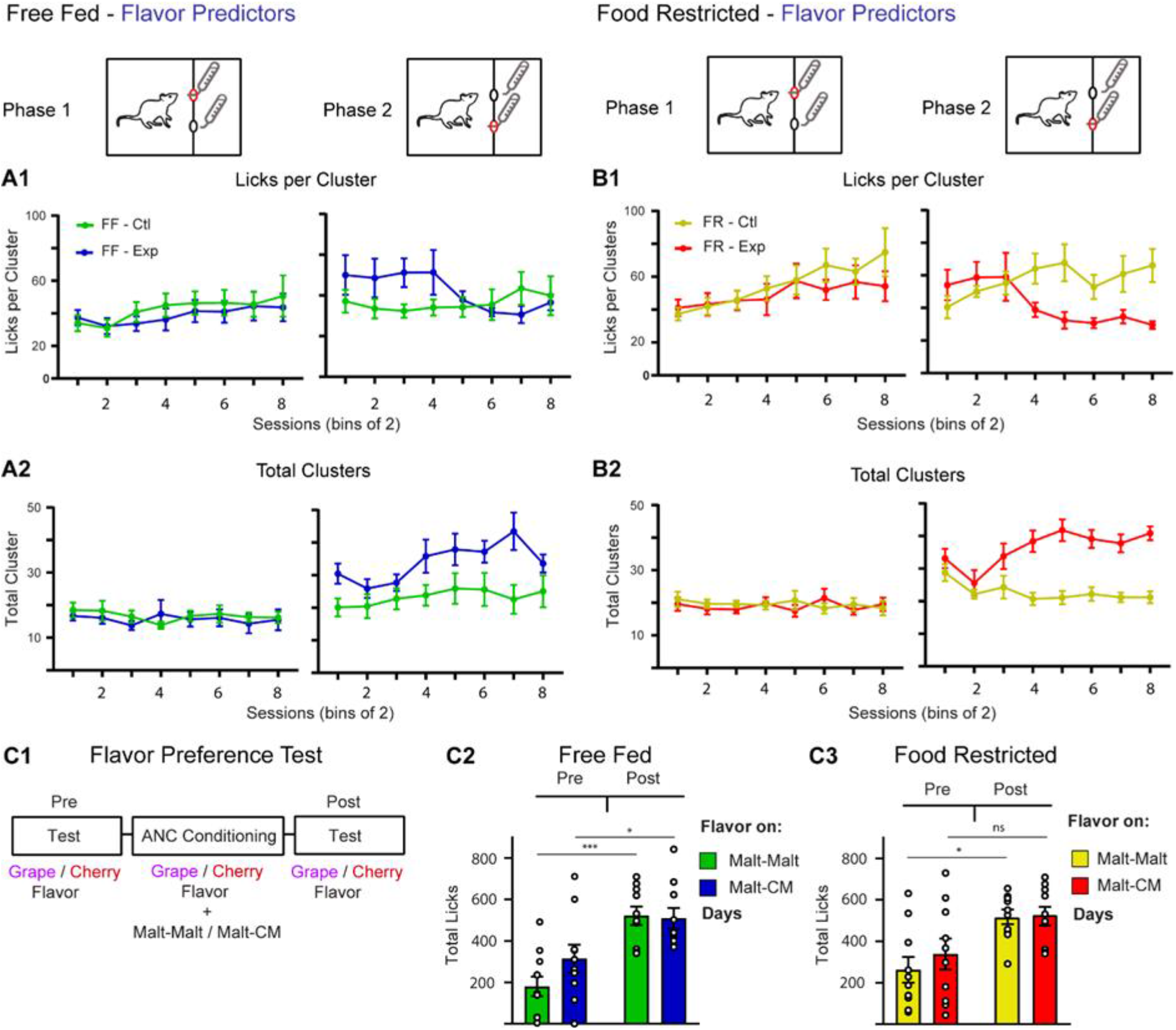
Lick microstructure analysis and flavor preference test for flavor contrast experiment. In both FF **(A1)** and FR **(B1)** animals, there were no differences phase 1 licks per cluster (left) between Malt-Malt and Malt-CM sessions while in phase 2. For total cluster bouts there was no difference between Malt-Malt and Malt-CM days for both FF **(A2)** and FR **(B2)** in phase 1 (left) while in phase 2 (right) there was a progressive increase in lick cluster bouts in Malt-CM compared to Malt-Malt sessions for both FF and FR animals. **C1**. Outline of flavor preference test and pre/post total lick responses for both flavors in FF **(C2)** and FR **(C3)** rats. FF, free-fed; FR, food restricted.

The amount of total clusters in phase 1 for either FF or FR rats did not differ in Malt-Malt compared to Malt-CM sessions (FF, 2-way ANOVA, Condition: F(1,9)=0.90, P=0.36; Session: F(7,63)=0.41, P=0.89; Interaction: F(7,63)=0.51, P=0.82) (FR, 2-way ANOVA, Condition: F(1,9)=0.25, P=0.62; Session: F(7,63)=0.22, P=0.98; Interaction: F(7, 63)=1.06, P=0.40) indicating that changes in total clusters does not contribute to changes in phase 1 total licks. There was however a significant increase in phase 2 total clusters in response to CM for both FF and FR animals throughout conditioning (FF, 2-way ANOVA, Condition: F(1,9)=11.73, P=0.008; Sessions: F(7,63)=2.90, P=0.01; Bonferroni post-hoc differences – Paired Sessions 4-7; No Interaction: F(7,63)=1.68, P=0.13) (FR, 2-way ANOVA, Condition: F(1,9)=31.07, P=0.0003; Session: F(7,63)=2.75, P=0.01; Interaction: F(7,63)=4.06, P=0.001; Bonferroni post-hoc differences – Sessions 4-8) which is in contrast to the changes in lick microstructure for contextually-driven ANC. These differences in the lick microstructure between animal using contextual (Figure 4) and gustatory predictors (Figure 7) suggest that the motivational processes underlying ANC for each are distinct.

#### Flavor preference tests show an increase lick frequency to both flavors following conditioning

To shed light on the hedonic changes in reward properties that underlie ANC we next tested whether the flavor cues without maltodextrin gain appetitive value as a result of contrast conditioning using a flavor + saccharin preference test performed. Grape and cherry flavored 0.2% saccharin solution without Malt or CM were made available before and after ANC conditioning to determine baseline flavor preference and changes in preference as a result of conditioning (Figure 7C1). For FF rats, a 2-way repeated measures ANOVA revealed a significant effect by test (pre vs post conditioning) but no main effect by Condition or Interaction between the two (Pre/Post: F(1,9)=58.04, P<0.0001; Condition: F(1,9)=2.12, P=0.18; Interaction: F(1,9)=1.83, P=0.21) indicating that there was no initial preference before conditioning and that licking to both flavors, irrespective of which flavor predicted CM, increased after conditioning (Figure 7C2). Similar results were seen for FR rats, with the exception that the main effect by Pre/Post was mostly driven by the flavor given on Malt-Malt days (2-way ANOVA, Pre/Post: F(1,9)=24.21, P=0.0008; Condition: F(1,9)=0.34, P=0.56; Interaction: F(1,9)=0.35, P=0.57) (Figure 7C3). These results suggest that pairing flavors with Malt enhances intake of both gustatory cues, regardless of which one predicts CM.

#### Contextual predictors are more effective ANC cues than gustatory predictors

In the current study, four unique contrast conditions were tested (Context: FF and FR; Flavor: FF and FR). To formally compare each condition, we next analyzed the normalized phase 1 total lick data (= total licks: Malt-CM/Malt-Malt) to determine the effectiveness of each condition compared to one another in promoting ANC (Figure 8). A 3-way ANOVA used to compare the results by Diet, Predictor, and Session showed a main effect by Session and an Interaction between Session and Predictor (Session: F(3.46,124.6) = 6.57, P=0.0002; Session X Predictor: F (7, 252) = 2.07, P=0.04; No effect of Diet: F(1, 36)=0.3132, P=0.58) indicating that, compared to flavor cues, the contextual predictors used here are slightly more effective in promoting ANC for Malt-CM rewards.

**Figure 8:**
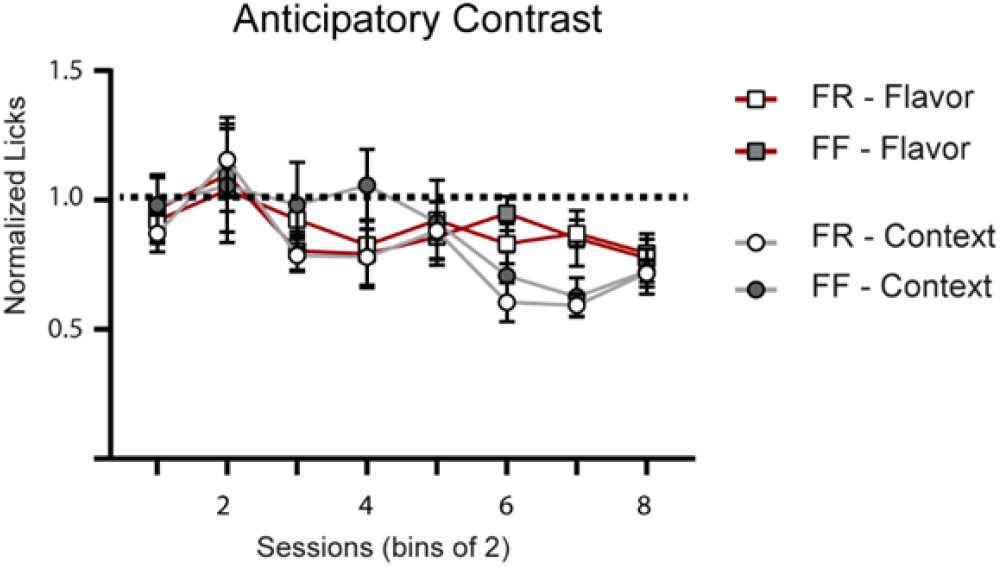
A comparison of all four ANC groups showing a significant difference in the effectiveness of contextual over gustatory predictors for ANC development. Data was normalized by dividing total licks on experimental day by total licks on the paired control day. FF, free-fed; FR, food restricted.

## Discussion

In the current study, we tested the predictive ability of contextual and gustatory cues on ANC using a novel sequence of nutritionally-relevant rewards and the motivational processes that underlie this behavior. Our results indicate that both contextual and gustatory information are effective predictors for ANC to develop. The selective reduction of Malt consumption is unlikely due to spatial competition between the two sipper ports as increases in premature port entries in phase 1 mostly preceded changes in total licks and magnitudes did not closely match ANC levels. In contextually-driven anticipatory sessions, our lick microstructure analyses suggest that ANC may be determined by a change in the rewarding properties of the Malt solution that is in all other respects equivalent to the one presented on control days - the only difference being the context signaling the availability of CM in phase 2. In contrast, ANC driven by gustatory predictors was not mirrored by changes in the lick microstructure indicating that the contrasting effects seen here are perhaps driven by competing motivational factors. Due to their biological relevance, gustatory cues can be robustly associated with food sources, potentially acting as strong secondary reinforcers that can compete with negative contrast effects (Lucas and Timberlake, 1992; Flaherty et al. 1995). Since the Malt solution is itself rewarding (Sclafani, 1987), animals may develop a strong liking for both flavors, irrespective of whether or not one also predicts CM in phase 2. In line with this idea, our flavor preference tests demonstrate a statistically equivalent increase in licking for both flavors after ANC conditioning.

Stereotypical patterns of rodent licking behavior are thought to reflect different underlying motivational processes where the amount of licks in a cluster of continuous licks has been shown to relate to reward palatability (reward liking) while the amount of total clusters changes with incentive value (reward wanting) as well as signal its post-ingestive feedback properties (Davis and Smith, 1992; reviewed in Dwyer, 2012; Johnson, 2018 and Naneix et al., 2019). Support for these parameters as useful metrics of reward properties has come from studies investigating changes in the hedonic characteristics of rewarding solutions where alterations in lick dynamics have been shown to correspond to changes in reward magnitude (Davis and Perez; 1993) or the motivational state of the animal (D’Aquila et al. 2012; Murphy et al., 2018). In the present study, only contextually-driven ANC showed a corresponding change in the lick microstructure where a change in licks per cluster was seen to develop between control and experimental days as ANC developed. These results are consistent with recent work by Wright et al., (2013) showing differences in licks per cluster that were related to contextually-driven ANC using sucrose rewards of different magnitude. In phase 2, CM lick patterning was also different in the two predictor groups despite both groups showing similar total CM lick rates. In both free-fed and food restricted groups, licks per cluster and total clusters for CM increased in sessions predicted by contextual cues while for animals relying on gustatory cues there was shift from a licks per cluster strategy to more licking bouts as ANC developed. Interestingly, compared to contextual cues, gustatory predictors for food restricted rats resulted in more total licks in phase 1 (Total licks: Context, 11567 ± 874; Flavor, 14968 ± 915 Unpaired T-Test, P=0.02) suggesting an increase in the incentive value of the flavored Malt solutions – an interpretation which is consistent with the results of the flavor preference test. Thus, it is possible that the addition of a flavor to the maltodextrin solution in phase 1 enhances its potential palatability vs. a non-flavored maltodextrin solution. This change in Malt palatability may in effect be impacting the hedonic disparity between the two rewards by altering the incentive value of CM. Such changes are reminiscent of those seen in temporal discounting paradigms where a comparative analysis of two rewards separated by time can impact the incentive value of each (Loewenstein, G., 1987; Myerson and Green, 1995; Cardinal, 2006). Despite showing comparable total CM lick rates, our data show that, with experience, differences in CM lick patterning appear between gustatory and context groups suggesting that the predictive and hedonic value of the phase 1 solutions are differentially affecting the motivational processes underlying CM consumption.

Occasion-setters have been shown to be at play when features are trained so that they disambiguate the relationship of another stimulus with an outcome (Holland, 1992). As such, the within-subjects nature of the current design using different predictors warrants a description of the paradigm in terms of occasion setting mechanisms. The target Malt is followed by the rich CM on half of the occasions, and therefore is an ambiguous predictor of CM. It is under such ambiguity that the context (Experiment 1) and the flavor (Experiment 2) stimuli disambiguate the meaning of Malt. In other words, Malt is followed by CM only when a feature is present. This design is thus reminiscent of an occasion setting design. One characteristic of occasion setting is that it occurs best when the occasion setter is presented serially with the target stimulus that it disambiguates. Although occasion setting has been observed with both simultaneous and serial compounds, it is much stronger with serial compounds (Holland, 1992; Fraser & Holland, 2019). This parallels the present results in that better ANC was observed when contextual cues were used relative to when flavor cues were used. Because contextual cues were experienced before Malt, whereas flavor cues were experienced simultaneously with Malt, the advantage of serial over simultaneous presentation of stimuli seen in occasion setting experiments can explain the difference between contextual and flavor cues observed in the current experiments.

The fact that contrasting effects are based on relative rather than the absolute value of the rewards make them sensitive to various factors. For instance, their value-based as well as temporal disparity can strongly impact contrast (Flaherty et al., 1994; Flaherty and Mitchell, 1999; Moss et al., 2002; Cottone et al., 2008). The motivational state of the animal is another key factor and experiments have shown that food deprivation can result in animals not developing ANC or even showing positive induction where animals increase consumption to the less rewarding option (Flaherty et al., 1991; Weatherly et al., 2005). In the current study, all food-restricted animals developed ANC. For those using contextual cues as predictors, these differences occurred after fewer conditioning trials than free-fed animals. Moreover, these animals consumed more CM than free-fed animals demonstrating a preferential food seeking approach for maximizing CM intake (CM Total licks: FR, 20817 ± 963.0; FF, 17500 ± 859 Unpaired T-Test, P=0.02 (data not shown)). These results may be attributed to the novel food choice sequence (Malt followed by CM solution) where the disparity in the hedonic value and nutritional content between the two was different than past investigations (Lucas et aI., 1990; Flaherty and Mitchell, 1999; Cottone et al., 2008; Parylak et al., 2012). Condensed milk has been shown to be a strong reinforcer as reward seeking studies have shown similar effort-based responding for condensed milk and cocaine (Ciccocioppo et al., 2004; Martin-Fardon et al., 2016; Schmeichel et al, 2018). This highly palatable food choice may thus be effective in promoting a selective feeding strategy even when animals are food restricted. The current findings thus indicate that, despite the benefit in taking an opportunistic approach to maximally consume in both phases, food restricted animals can act selectively in their feeding choices so as to potentially maximize their intake of a nutritionally-rich food option, and that learning mechanisms underlie these choices.

A number of different neural process must be at play when animals learn to predict rewards of different magnitude. The lick microstructure analysis performed in the current study suggests that gustatory and contextual predictors may be mediating ANC through different motivational processes. Studies investigating the neurobiological mechanisms that underlie ANC have mostly focused on the changes in reward value that might contribute to ANC development. While dopamine activity encodes information predicting a future reward as well as the reward itself (Bromberg-Martin and Hikosaka, 2010) the role it plays in ANC development is unclear. Using systemic administration of the monoamine stabilizer (−)-OSU6162, Feltmann et al., (2018) showed that it had no impact on anticipatory contrast. Moreover, lesions of the nucleus accumbens (NAc) – a region important for integrating dopamine-based reward processing – have no effect on ANC (Leszczuk and Flaherty, 2000) suggesting that alternative regions may be involved. One candidate circuit might be prefrontal cortex (PFC) where reward based and sensory signals converge for supporting memory (Groenewegen and Uylings, 2000). In addition, PFC functioning has been heavily implicated in cognitive control mechanisms for selecting appropriate actions (Miller, 2000) often with delays imposed between stimulus and response (Mobini et al., 2002; Kable and Glimcher, 2007; Kin et al., 2009). As ANC requires the integration and working memory representation of sensory content predicting reward the PFC may play an important role in ANC development.

In summary, our results indicate that despite the potential benefit in taking an opportunistic feeding approach, food-restricted animals can use contextual predictors to act selectively in their feeding choice and that these changes may stem from learned changes in the hedonic properties of the readily available food source. Gustatory predictors can also be used to optimize intake of preferred food option in both free-fed and food restricted rats but less effectively than contextual cues - a results that may be due to the high predictive strength of flavors linked to a food source that differentially impacts the motivational processes that drive ANC. Future investigation of the neural activity contributing to these motivational changes will help elucidate the neurobiology that allows animals to optimize their foraging strategies.

## Acknowledgements

The authors acknowledge the help and support from the staff of the Division of Biomedical Services, Preclinical Research Facility, University of Leicester, for technical support and the care of experimental animals as well as colleagues in the department of neuroscience, psychology and behavior at the University of Leicester for their academic contribution. Also thanks to Charlotte Bonardi for helpful discussions on the occasion setting. This work was funded by the Wellcome [grant #209023/Z/17/Z to J.A-S] and the Leverhulme Trust [grant #RPG-2017-417 to J.E.M. and J.A-S.].

